# Missing kinaesthesia challenges precise naturalistic cortical prosthetic control

**DOI:** 10.1101/004861

**Authors:** Ferran Galán, Mark R. Baker, Kai Alter, Stuart N. Baker

## Abstract

A major assumption of brain-machine interface (BMI) research is that patients with disconnected neural pathways can still volitionally recall precise motor commands that could be decoded for naturalistic prosthetic control. However, the disconnected condition of these patients also blocks kinaesthetic feedback from the periphery, which has been shown to regulate centrally generated output responsible for accurate motor control. Here we tested how well motor commands are generated in the absence of kinaesthetic feedback by decoding hand movements from human scalp electroencephalography (EEG) in three conditions: unimpaired movement, imagined movement, and movement attempted during temporary disconnection of peripheral afferent and efferent nerves by ischemic nerve block. Our results suggest that the recall of cortical motor commands is impoverished in absence of kinaesthetic feedback, challenging the possibility of precise naturalistic cortical prosthetic control.

---

Brain-machine interface (BMI) technology offered early promise of restoring independence to those with spinal cord injury. Grounded in seminal work from awake behaving monkey^1,2^, BMI research aims to decode movement parameters from neural ensemble activity, enabling natural control of assistive devices. However, the speed and accuracy of current BMIs^3,4^ are poor compared to natural movements and highly dependent on visual feedback, showing some similarities with motor deficits in patients with sensory neuropathies. This observation led us to speculate that absent^3,4^ and arbitrary^5^ sensory feedback produced by controlling an artificial actuator (e.g. robotic arm) interferes with the recruitment of the neural population previously engaged in controlling a natural effector with intact feedback. Rather than viewing the spatiotemporal sequence of activity which produces movement as internally generated by cortical circuits in a feedforward manner^6,7^, such a view would extend the ‘dynamical machine’^7^ responsible for movement to include afferent feedback from the periphery. This is supported by evidence for the rapid integration of sensory feedback into motor output, whilst taking account of high-level movement goals^8,9^. In such a framework, loss of feedback would have an impact comparable to the lesion of a cortical area. Movement might still be possible, but only after reconfiguration of the network, and is likely to be impoverished compared with the natural state. It is known that primary sensory and motor areas undergo plastic changes^10-13^ associated with abnormal function^14^ when afferent inputs are removed. Moreover, studies with amputees suggest that access to the motor representation of the missing limb is conditional upon the re-establishment of peripheral connections and restoration of the sensorimotor loop^15^, an observation also supported by experiments in patients with hand allografts^16^ and targeted muscle re-innervation for prosthetic control^17^.

We hypothesized that cortical motor commands cannot be effectively generated in the absence of kinesthetic feedback. Here we tested this hypothesis, using temporary ischemic nerve block to model disconnection of cortical circuits from the periphery. In the same subjects, we compared EEG decoding of unimpaired movements (*Move/MoveAfter)* with the same movements attempted during peripheral disconnection (*Block;* see Fig. 1a). We further evaluated decoding of imagined movements (*Imag*), which also lack movement re-afference and have been utilized to calibrate decoders for people with tetraplegia^4,18^ (Fig. 1a, b). We found that effective decoding was only possible when displacement-triggered re-afference was present, suggesting that cortical motor commands deteriorate when they cannot be updated by their sensory consequences. This challenges the possibility of precise naturalistic cortical prosthetic control in patient groups with peripheral disconnection.

**Figure 1.**
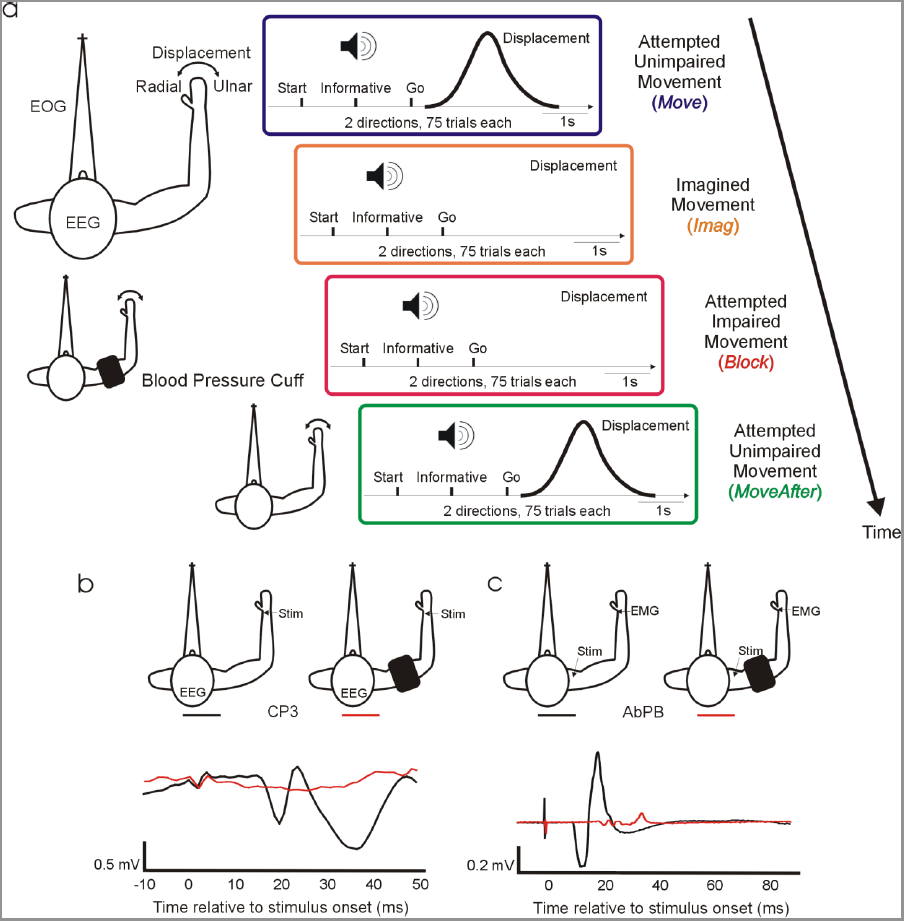
Experimental Setup. **(a)** Behavioral task. Diagrams depict the temporal structure within trials and between conditions. Auditory cues indicate trial start, instructed deviation, and ‘go’. Schematic views of the arm, from above, describe subjects’ position and experimental setup during task performance. The top subject schematic, corresponding to *Move* and *Imag*, displays the electrophysiological signals and displacement measured during all experimental conditions. The middle subject schematic illustrates the position of the blood pressure cuff during *Block* condition. The bottom subject schematic corresponds to *MoveAfter*. Only conditions *Move* and *MoveAfter* involve movements. (b) Afferent block. Example of averaged SEP recorded at CP3 following median nerve stimulation before (black) and after (red) deafferentation by the ischemia. After deafferentation N20 is absent. (c) Efferent block. Example of CMAP recorded at muscle abductor pollicis brevis following supramaximal Erb’s point single pulse magnetic stimulation before (black) and after (red) motor block

## Results

Source analysis revealed significant differences between conditions in the dynamics of the cortical sources generating movement related EEG scalp potentials (Fig. 2a, b). We defined regions of interest (ROIs) to encompass pre- and post-central cortex based on standard Atlas coordinates (Tzourio-Mazoyer), and measured mean source absolute activation over these regions (Fig. 2c). We also defined different time windows within a trial, and measured mean source absolute activation over these windows (Fig. 2a, d). Activity in the pre-central ROI was significantly increased (p<0.01) in *Move* condition compared with *Imag* and *Block* during movement preparation (window 6), and significantly increased (p<0.01) in *Move* and *MoveAfter* compared with *Imag* and *Block* during maximum displacement (window 10); for a post-central ROI, significant differences were also seen in *Move* and *MoveAfter* compared with *Imag* and *Block* during maximum displacement (window 10). Neither pre-central nor post-central ROI activity differed significantly between conditions during any other analyzed window (see Fig. 2d-e).

**Figure 2.**
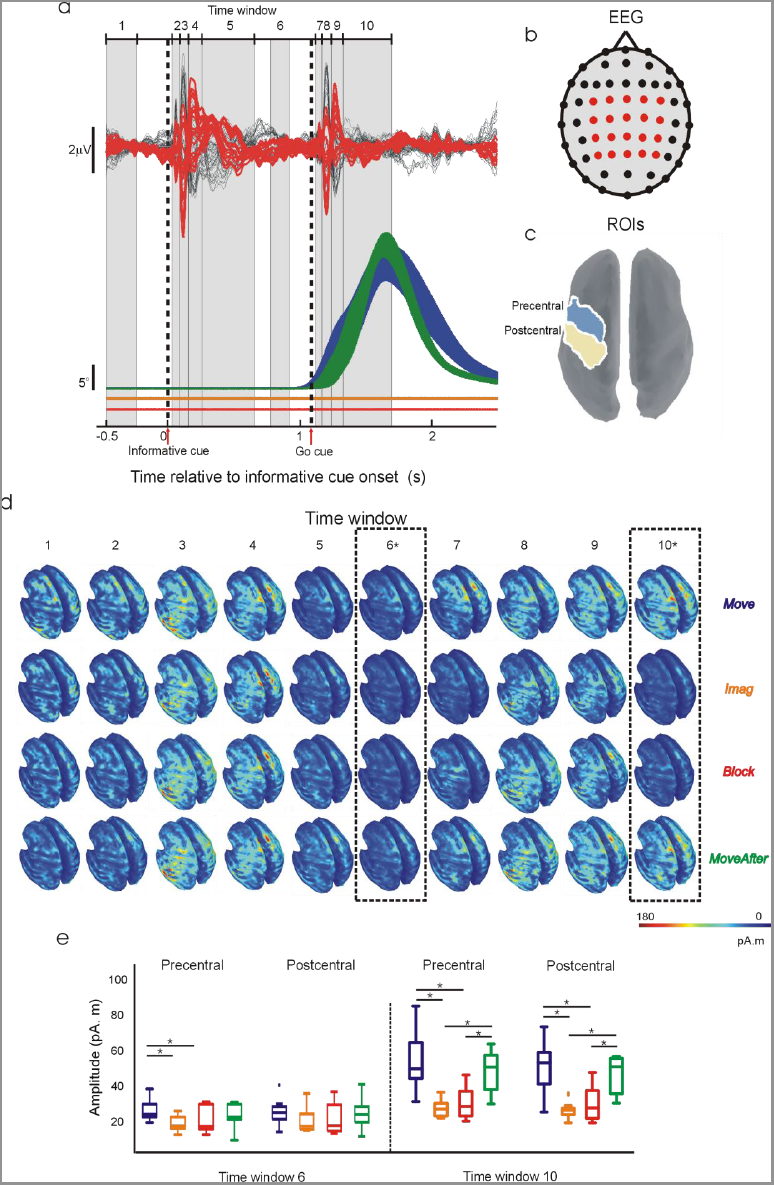
Temporal dynamics of EEG source activity. (a) Top, EEG grand averages across subjects. In red, traces corresponding to channels included in decoding analysis. Bottom, averaged (*±SEM*) absolute displacement across subjects. Color codes condition as in Fig. 1a. Displacement traces for *Imag* and *Block* conditions have been shifted downwards slightly for clarity. Vertical dashed lines represent ‘informative’ and ‘go’ cue onsets. Grey numbered boxes define the 10 windows utilized for analysing the dynamics of EEG source activity (see Methods). (b) EEG channels’ topographic distribution. Red channels correspond to red traces in **a. (c)** Localization of pre-central and post-central ROIs utilized for analysing EEG source activity. **(d)** Grand averages across subjects of time averaged absolute cortical source activity within each time window defined in a. Dashed line boxes indicate windows in which significant differences were found between conditions. **(e)** Box plots of averaged source activity at pre-central and post-central ROIs from each subject in time windows that revealed significant differences between conditions (windows 6 and 10 highlighted with dashed 2line boxes in **d)**. * In **d** and **e** denotes significant difference (p<0.01).

The activity of the pre-central sources responsible for generating motor output was diminished in the absence of kinaesthetic feedback; however, this does not necessarily mean that the underlying motor commands which this reflected were impaired. To probe this in more detail, we carried out a decoding analysis which attempted to predict the movement direction based on the EEG. We reasoned that if the smaller signals seem in *Imag* and *Block* conditions were still capable of good movement decoding, this would indicate some motor command nevertheless remained intact. In *Move* and *MoveAfter* conditions decoding performed significantly better (69.4% ± 3.1% and 66.3% ± 2.9% respectively, p<0.01) than by chance, but this was not the case for *Block* and *Imag* (51.6% ± 0.6% and 54.2% ± 1.3% respectively, significantly lower than *Move* and *MoveAfter*, p<0.01). Importantly, the plots of time-resolved decoding accuracy (Fig. 3a) revealed that peak accuracy in the *Move* or *MoveAfter* conditions was temporally confined around task performance (see tick marks on and below color maps in Fig. 3a), as expected for a genuine neural signal. By contrast, peak decoding in *Imag* or *Block* conditions was distributed randomly over the analyzed timeframe, suggesting chance decoding of noise fluctuations unrelated to underlying neural processes.

**Figure 3.**
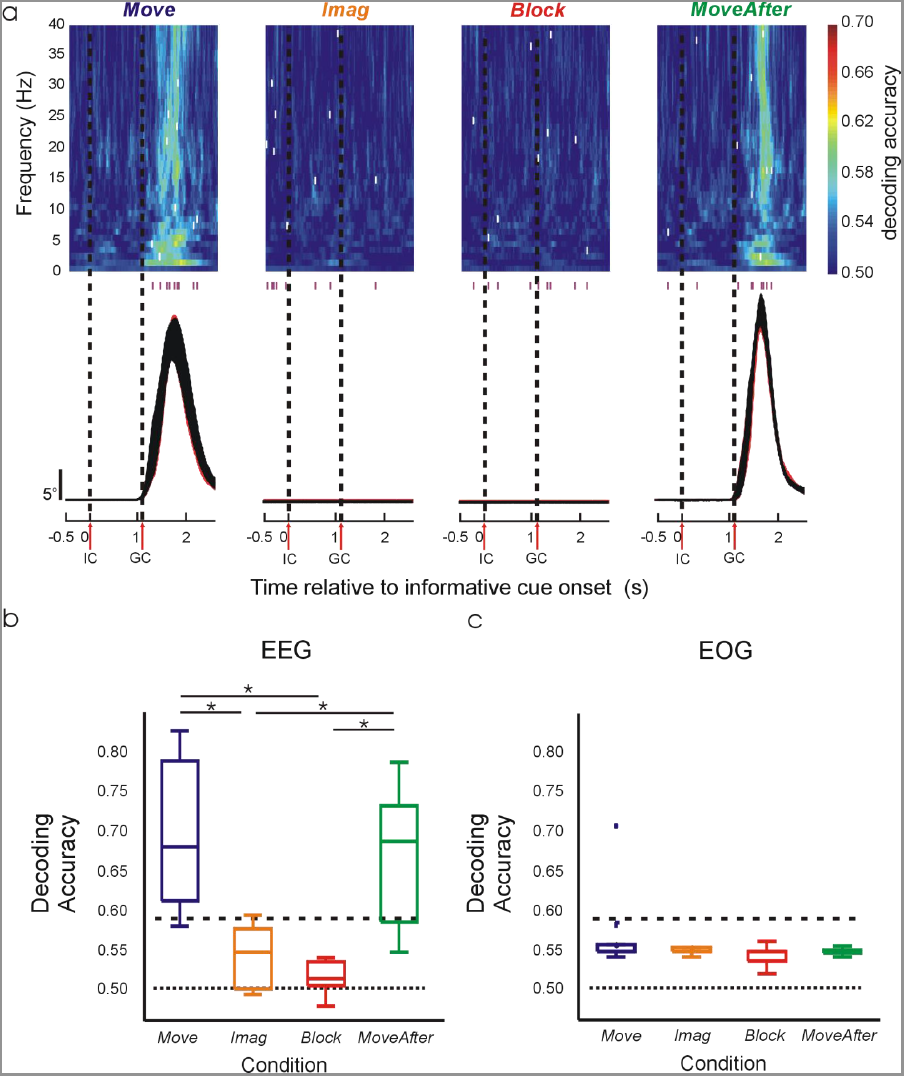
Decoding of wrist deviation. (a) Top, subject-averaged decoding accuracy (DA) in time-frequency space using EEG signals. White ticks within the color map indicate time-frequency bins with maximum DA from each subject; the time of these bins is also shown with purple ticks beneath the color map. Bottom, average absolute displacement (*±SEM*) across subjects. Vertical dashed lines represent ‘informative’ and ‘go’ cue onsets. (b) Box plots of DA from each subject using EEG signals. (c) Box plots of maximum DA from each subject using EOG signals. Horizontal dotted and dashed lines represent chance level of 50% and the value above which DA deviates significantly (p<0.05) from the chance level. *Significant pairwise difference (p<0.01).

Further evidence of a key difference in the nature of the signals decoded came from source analysis at the time sample of maximum decoding accuracy (Fig. 4a). As expected, in the *Move* condition the contralateral primary sensorimotor areas were the major contributors of EEG surface activity. Activity in the pre-central and post-central ROI was significantly reduced (p<0.01) in *Imag* and *Block* conditions compared with *Move* (Fig. 4b). Activity in *MoveAfter* seemed to restore partially, although not completely, back to that seen in *Move* (see Figs. 2e, 3b, 4b).

**Figure 4.**
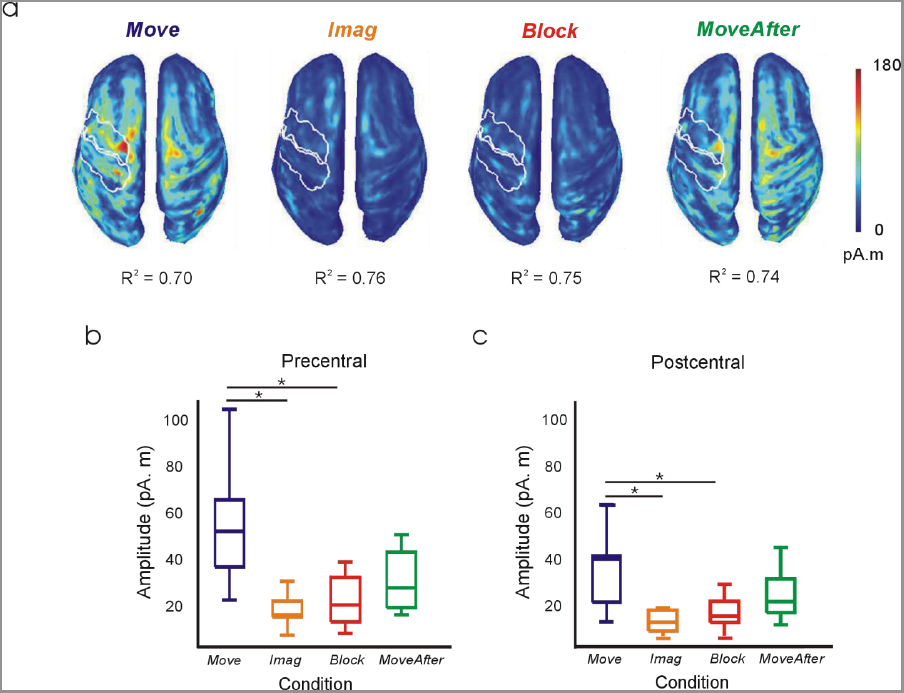
Cortical source activity during maximum DA. **(a)** Subject-averaged absolute value of cortical source activity at time bins of maximum DA (see white ticks in Fig. 3a). Spatial goodness of fit of source activity in each condition is indicated by averaged R^2^ values. White outlines define contralateral pre-central and post-central ROIs. **(b, c)** Box plots of averaged source activity at pre-central and post-central ROIs from each subject. *Significant pairwise difference (p<0.01).

One possible confound with EEG decoding is that scalp potentials could be subtly influenced by eye movement artifacts. To check for this, we carried out a similar decoding analysis as above, but using EOG signals (Fig. 3c). No significant decoding occurred in any condition, indicating that eye movements were uncorrelated with the instructed wrist movements and that EOG contamination of the EEG could not explain our results.

## Discussion

This study shows that movement-related information contained in EEG is impoverished in the absence of kinaesthetic feedback.

Previous work has correlated various movement parameters with motor cortical activity ^1,2,19-29^, establishing the scientific basis for developing BMIs^30-35^ that could confer intuitive neuroprosthetic control to patients suffering from paralysis^3,4^. However, experiments with patients suffering large fiber sensory neuropathies^36-39^ have shown that without visual feedback the motor output of these patients fluctuates randomly. Kinaesthetic feedback plays an important role in motor control by allowing for error-correction and by contributing in the formation of accurate internal models of limb dynamics^40,41^. These observations pose a relevant question to BMI research that, to our knowledge, has been unaddressed: to what extend does missing kinaesthesia prevent the generation of the normal sequence of motor commands required for voluntary movements?

Here, we addressed this question by decoding from EEG activity attempted movements impaired by nerve block and imagined movements that lack movement re-afference. Both conditions are relevant to our goal, but have their limitations. Ischemic nerve block provides a well-established reversible model of short-term amputation-induced cortical reorganization in humans^10,15^ and provides a valid model for the loss of feedback from large diameter sensory fibers. However, the rapid onset of feedback loss, including the tonic level of drive seen in the steady state, could produce acute changes in cortical function not directly related to loss of movement-related feedback. On the other hand, imagined movements lack movement re-afference, but preserve tonic feedback about the (unchanging) limb state. In imagined movements, central mechanisms presumably are also acting to prevent overt motor outflow^42^. Importantly, imagined movements have been utilized to calibrate decoders for people with tetraplegia^4,18^. By using both of these non-invasive techniques in healthy subjects we were able to compare, for the first time, decoding of unimpaired, impaired and imagined movements. This overcomes some of the limitations presented by experiments with paralyzed patients and non-human primates: paralyzed patients are unable to perform goal-directed movements (an essential condition to which all others should ideally be compared); by contrast, it is virtually impossible to control for non-task related movements and to assess the quality of motor imagery in non-human primates.

Our study revealed that the lack of re-afferent feedback in imagined and impaired movements had an impact on the dynamics of the cortical sources generating movement related EEG scalp potentials. This appeared as reduced pre-central cortical activity during preparation, and reduced pre-central and post-central cortical activity during execution of movements. Three further observations support the notion that cortical activity became grossly abnormal in the absence of sensory feedback. First, decoding accuracy dropped to chance levels. Second, the time of maximal decoding accuracy was no longer temporally locked around the time of attempted movement. Third, pre- and post-central cortical activity at the time of maximum decoding was substantially reduced. Our findings extend to deafferentation those by other studies reporting reduced activity^43,44^ and reduced decoding or tuning^44,45^ when comparing imagined with unimpaired movements, highlighting the critical role of kinaesthetic feedback for the successful recruitment of the neural population responsible for precise movement control.

Our study used EEG to access the neural signals underlying motor commands, and showed that decoding efficiency was reduced in the absence of a normal sensorimotor loop. We cannot exclude the possibility that motor processing continued relatively intact in these tasks, but that only the overt manifestation as discriminable scalp potentials was degraded. If so, this would imply that only EEG-based naturalistic cortical prosthetic control will be challenged in paralyzed patients lacking feedback, whereas BMIs relying on invasively recorded single unit activity may still operate effectively. However, several pieces of evidence argue that our findings may also have applicability to invasive BMI. Pandarinath and colleagues^45^ have reported weakened movement-related multiunit modulation in human primary motor cortex (M1) during imagined movements, demonstrating that such degradation is also present at the neural scale. Other studies have demonstrated the influence of kinesthetic feedback in ongoing M1 activity^8,46-48^ suggesting the importance of trans-cortical feedback pathways for predicting optimal states^49^ and for modulating sensory feedback accordingly^50^.

Non-biomimetic approaches, which require the subject to learn to modulate arbitrary but readily discriminable signatures of neural activity^51,52^, represent a different approach to BMI which has been explored in paralyzed patients^53^. However, it is an open question whether the rate with which subjects can learn to use such systems, or the eventual performance obtained, is also affected by the lack of a functional sensorimotor feedback loop.

Cole^54^ provides a poignant description of the devastating consequences for one patient which sudden loss of large-diameter afferents had for motor control. Although voluntary movement was regained after much retraining, this required great concentration, and strongly depended on visual feedback. Current cortical prosthetic control resembles Cole’s description. Future research is needed to explore alternatives to compensate for such kinaesthetic loss^5^, which would allow intuitive control of prosthetic devices while maintaining an intact sensorimotor feedback loop.

## Methods

### Experimental Setup

Nine healthy adults (7 men, 2 women) took part in the study. All the experimental procedures were approved by the Research Ethics Committee of the Medical Faculty, Newcastle University; subjects provided written informed consent to participate. Each subject performed four experimental conditions in the following order: unimpaired movement (*Move*), imagined movement (*Imag*), attempted movement after ischemic nerve block (*Block*) and unimpaired movement after circulation had been restored to the arm (*MoveAfter*) (see Fig. 1a). Subjects sat comfortably in a chair fixating on a static visual target placed approximately 150 cm ahead. The right elbow and pronated forearm were gently placed at waist height onto an arm rest, which restricted movement to radial and ulnar deviation at the wrist joint (lateral hand movements). The hand was placed flat with extended fingers on the rest, which was padded with memory foam. The same foam imprints were used across sessions to minimize postural changes. The rest was custom made to fit each subject’s arm well, minimizing hand movements and movement around other joints. Wrist angular displacement was sensed by a potentiometer, fixed with its axis coaxial to the wrist joint. A displacement of 0° indicated the neutral position with the hand in the same plane as the forearm; positive angles denoted radial deviation. The left arm rested unrestrained in a comfortable position throughout the task. Each trial commenced with an auditory start cue (2000 Hz; 100 ms), followed 1s later by an informative cue (100 ms) indicating the required direction on that trial: radial (1500 Hz) or ulnar (500 Hz) deviation. After a 1s delay period a ‘Go’ cue (1000 Hz; 100 ms) indicated when to initiate the movement. In *Move* and *MoveAfter* conditions subjects were instructed to perform fast, stereotyped radial/ulnar deviations of the wrist. In the *Imag* condition subjects were requested to imagine radial/ulnar deviations as performed during the immediately-preceding *Move* trials but without overt movement. In *Block* condition subjects were instructed to try to perform the movements as in *Move*, despite the impairment produced by nerve block. Each condition consisted of a randomized sequence of 150 trials (75 in each direction) and lasted 13 minutes.

### Data Acquisition

Scalp EEG was recorded by a 61–sensor cap according to the International 10–20 system referenced to Cz. Electro-oculograms (EOG) were recorded via bipolar electrodes; horizontal EOG was recorded by placing an electrode to the outer canthus of each eye, and vertical EOG by an electrode pair above and below the subjects’ left eye. Both EEG and EOG signals were sampled at 1 kHz (Neuroscan SynAmps 2RT, Compumedics USA, Charlotte, NC) and grounded with an electrode placed over the left clavicle. Impedance for all electrodes was <5 kΩ. To record stimulus-evoked responses (see below), bipolar surface electromyogram (EMG) was recorded from the right abductor pollicis brevis (AbPB), amplified and high-pass filtered at 30 Hz (D360, Digitimer, Welwyn, UK), and sampled at 5 kHz (CED Micro1401, Cambridge, UK). A ground electrode was placed on the dorsum of the wrist.

### Ischemic Nerve Block Procedure

Ischemic nerve block was achieved by applying a commercial blood pressure cuff to the right arm at the level of the biceps. The arm was first raised for ∼30s to drain blood from the large veins; the cuff was then rapidly inflated to a pressure of 180 mm Hg. The arm was then gently lowered and placed into the apparatus. Cuff pressure was maintained constant through this experimental condition. Subjects were told not to contract muscles distal to the cuff, from the moment that the arm was raised, until the experimental recording began. Because we emphasized the importance of this instruction, subjects were able to remain relaxed throughout, and thereby avoided muscle pain associated with lactate buildup.

Somatosensory evoked potentials (SEPs) following electrical stimulation of the median nerve at the wrist (stimulus rate 9 Hz, pulse width 1 ms, intensity just below motor threshold, 1000 stimuli) and compound muscle action potentials (CMAPs) of the AbPB muscle following magnetic nerve stimulation (Magstim 200, Dyfed, UK; single pulse; intensity supra-maximal) at the supra-clavicular fossa (Erb’s point) were monitored to assess sensory and motor block.

Baseline SEPs and CMAPs were measured before applying the cuff. Beginning 18 minutes after cuff inflation, SEPs were measured at intervals of two minutes until the contralateral N20 was absent, indicating complete block of large fiber sensory afferents (see Fig. 1b). At this stage the subjects no longer perceived the nerve stimulus. Two minutes after this point, CMAPs were measured at intervals of 30 s until their amplitude was significantly reduced, indicating almost complete motor block (see Fig. 1c). Subjects were then requested to attempt to perform the task (*Block* condition). Immediately after task completion (75 trials in each direction) the cuff was removed, limiting the maximum total ischemic time to 50 minutes. The *MoveAfter* condition started 5–10 minutes after cuff removal, when subjects no longer reported reperfusion paraesthesias.

### Data Analysis

#### Signal processing

EEG and EOG single trial epochs were extracted from 500 ms before to 2500 ms after the informative cue. In *Move* and *MoveAfter* conditions, trials with no movement, movement in the wrong direction, or movement with a reaction time five times the median absolute deviation above the mean were excluded. EEG data were filtered (1–40 Hz) and re-referenced (common average reference, CAR). Poor quality EEG channels were excluded before computing CAR. Time-resolved amplitudes of oscillations in the 1–40 Hz frequency range were computed using complex Morlet wavelets (2 s time resolution at 1 Hz central frequency). EOG data were low-pass filtered with a cut-off at 30 Hz (2nd order Butterworth, zero phase shifts).

#### Cortical sources of EEG surface activity

Cortical sources of single-trial EEG surface activity were estimated by computing Tikhonov-regularized minimum-norm estimates^55^ on a symmetric BEM head model^56^ using constrained dipoles (15000 vertices normal to cortical surface) and standard Tikhonov regularization (*λ* = 0.1). The cortical current maps were analyzed using regions of interest (ROIs) defined by Tzourio-Mazoyer atlas onto Colin 27 volume coordinates^57^. ROI activity was computed as the averaged absolute activity of all the vertices included in the ROI. Temporal dynamics of ROI activity was examined by measuring mean source activation over time samples included within 10 different time windows: −500 to −250 ms before informative cue onset (window 1), 15 to 60 ms (window 2), 60 to120 ms (window 3), 120 to 240 ms (window 4), 240 to 600 ms (window 5), 750 to 900 ms (window 6), 1115 to1165 ms (window 7), 1165 to 1220 ms (window 8), 1220 to1300 ms (window 9) and 1330 to 1700 ms after informative cue onset (window 10; see Fig. 2a, c).

Spatial goodness of fit of the estimated sources at the time bin of maximum DA was estimated with the averaged coefficient of determination (R^2^) across trials and subjects for each condition.

#### Decoding wrist deviation from electrophysiological signals

Wrist deviations were decoded by using a Bayes linear classifier (see classifier description below). Decoding accuracy in Fig. 3a was estimated for each condition by leave-one-out cross-validation across 40 frequency components and 3000 time bins (corresponding to 1–40 Hz range and 3 s epochs sampled at 1 kHz) using the estimated amplitudes from 20 EEG channels covering bilateral sensorimotor areas (FC3, FC1, FCz, FC2, FC4, C4, C2, Cz, C1, C3, CP3, CP1, CPz, CP2, CP4, P4, P2, Pz, P1, P3). Decoding accuracy in Fig. 3b was estimated for each condition by leave-one-out cross-validation combining through the arithmetic mean^58^ nine classifiers using the most informative time-frequency bin from each subject (bins indicated as white ticks in Fig. 3a). To assess whether wrist movement direction could be inferred from correlated eye movements, decoding accuracy was estimated for each condition by leave-one-out cross-validation across 3000 time bins (3 s epochs sampled at 1 kHz) using vertical and horizontal filtered EOG.

#### Linear classifier

A Bayes linear classifier^59^ was used to decode wrist deviation *d* ⊂ *J* = {*radial*, *ulnar*} from a signal vector of *N = 20* EEG or *N = 2 EOG*. The likelihood functions were modeled as multivariate Gaussian distributions according to

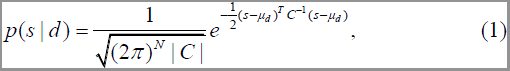

where *s* depicts either the 20 dimensional signal vector comprising the amplitudes of a single frequency component recorded from 20 EEG channels at a certain time bin, or the 2 dimensional signal vector comprising both filtered vertical and horizontal EOG at a certain time bin. *C* is the common, i.e. deviation independent, covariance matrix and *µ*_*d*_ the deviation specific mean signal vectors.

For classification, the posterior probabilities were computed using Bayes’ rule

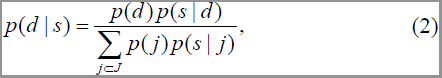

where a uniform prior with *p*(*d)* = 0.5 was used. Vector *s* was finally assigned to the deviation with the highest posterior probability.

#### Statistical analysis

Results for each condition are reported within the text as mean ± standard error of the mean (SEM). Within figures, the central line in box-plots is the median, the box is defined by 25th and 75th percentiles, the whiskers extend to the most extreme data points not considered outliers, and outliers are plotted individually (∼ ±2.7σ). Differences between the four experimental conditions were examined using Friedman’s test. Significance levels were corrected for multiple comparisons with a family-wise error rate *α* = 0.05. Significance of the decoding accuracy was examined with the cumulative binomial distribution^23^ using the lowest total number of trials across subjects (*n* =140) to obtain a statistically conservative significance.

All analysis was carried out offline in the MATLAB environment (The MathsWorks, Natick, MA, USA). EEG filtering, EEG re-referencing, EEG time-frequency decomposition and EEG source estimation were performed with Brainstorm^60^, which is documented and freely available for download online under the GNU general public license (http://neuroimage.usc.edu/brainstorm).

## Acknowledgements

This work was supported by the Wellcome Trust.

## Contributions

FG, MRB and SNB designed the study; FG, KA and SNB carried out the experiments; FG performed data analysis and wrote the first draft of the manuscript; all authors reviewed and critiqued the manuscript.

## References

1 Evarts, E. V. Relation of pyramidal tract activity to force exerted during voluntary movement. Journal of neurophysiology 31, 14–27 (1968).

2 Georgopoulos, A. P., Kalaska, J. F., Caminiti, R. & Massey, J. T. On the relations between the direction of two-dimensional arm movements and cell discharge in primate motor cortex. The Journal of neuroscience : the official journal of the Society for Neuroscience 2, 1527–1537 (1982).

3 Collinger, J. L. et al. High-performance neuroprosthetic control by an individual with tetraplegia. The Lancet 381, 557–564 (2013).

4 Hochberg, L. R. et al. Reach and grasp by people with tetraplegia using a neurally controlled robotic arm. Nature 485, 372–375, doi:10.1038/nature11076 (2012).

5 Weber, D. J., Friesen, R. & Miller, L. E. Interfacing the somatosensory system to restore touch and proprioception: essential considerations. Journal of motor behavior 44, 403–418, doi:10.1080/00222895.2012.735283 (2012).

6 Churchland, M. M. et al. Neural population dynamics during reaching. Nature 487, 51–56, doi:10.1038/nature11129 (2012).

7 Shenoy, K. V., Sahani, M. & Churchland, M. M. Cortical control of arm movements: a dynamical systems perspective. Annual review of neuroscience 36, 337–359, doi:10.1146/annurev-neuro-062111-150509 (2013).

8 Pruszynski, J. A. et al. Primary motor cortex underlies multi-joint integration for fast feedback control. Nature 478, 387–390, doi:10.1038/nature10436 (2011).

9 Krutky, M. A., Ravichandran, V. J., Trumbower, R. D. & Perreault, E. J. Interactions between limb and environmental mechanics influence stretch reflex sensitivity in the human arm. J Neurophysiol 103, 429–440, doi:10.1152/jn.00679.2009 (2010).

10 Brasil-Neto, J. P. et al. Rapid modulation of human cortical motor outputs following ischaemic nerve block. Brain : a journal of neurology 116 **(Pt** **3**), 511–525 (1993).

11 Cohen, L. G., Bandinelli, S., Findley, T. W. & Hallett, M. Motor reorganization after upper limb amputation in man. A study with focal magnetic stimulation. Brain : a journal of neurology 114 **(Pt** **1B**), 615–627 (1991).

12 Merzenich, M. M. et al. Topographic reorganization of somatosensory cortical areas 3b and 1 in adult monkeys following restricted deafferentation. Neuroscience 8, 33–55 (1983).

13 Sanes, J. N., Suner, S., Lando, J. F. & Donoghue, J. P. Rapid reorganization of adult rat motor cortex somatic representation patterns after motor nerve injury. Proceedings of the National Academy of Sciences of the United States of America 85, 2003–2007 (1988).

14 Cramer, S. C., Lastra, L., Lacourse, M. G. & Cohen, M. J. Brain motor system function after chronic, complete spinal cord injury. Brain : a journal of neurology 128, 2941–2950, doi:10.1093/brain/awh648 (2005).

15 Reilly, K. T., Mercier, C., Schieber, M. H. & Sirigu, A. Persistent hand motor commands in the amputees’ brain. Brain : a journal of neurology 129, 2211–2223, doi:10.1093/brain/awl154 (2006).

16 Vargas, C. D. et al. Re-emergence of hand-muscle representations in human motor cortex after hand allograft. Proceedings of the National Academy of Sciences of the United States of America 106, 7197–7202, doi:10.1073/pnas.0809614106 (2009).

17 Kuiken, T. A. et al. Targeted reinnervation for enhanced prosthetic arm function in a woman with a proximal amputation: a case study. Lancet 369, 371–380, doi:10.1016/s0140-6736(07)60193-7 (2007).

18 Hochberg, L. R. et al. Neuronal ensemble control of prosthetic devices by a human with tetraplegia. Nature 442, 164–171, doi:10.1038/nature04970 (2006).

19 Bradberry, T. J., Gentili, R. J. & Contreras-Vidal, J. L. Reconstructing Three-Dimensional Hand Movements from Noninvasive Electroencephalographic Signals. The Journal of Neuroscience 30, 3432–3437, doi:10.1523/jneurosci.6107-09.2010 (2010).

20 Bradberry, T. J., Rong, F. & Contreras-Vidal, J. L. Decoding center-out hand velocity from MEG signals during visuomotor adaptation. NeuroImage 47, 1691–1700, doi:http://dx.doi.org/10.1016/j.neuroimage.2009.06.023 (2009).

21 Jerbi, K. et al. Coherent neural representation of hand speed in humans revealed by MEG imaging. Proceedings of the National Academy of Sciences 104, 7676–7681, doi:10.1073/pnas.0609632104 (2007).

22 Kalaska, J. F., Cohen, D. A., Hyde, M. L. & Prud’homme, M. A comparison of movement direction-related versus load direction-related activity in primate motor cortex, using a two-dimensional reaching task. The Journal of neuroscience : the official journal of the Society for Neuroscience 9, 2080–2102 (1989).

23 Mehring, C. et al. Inference of hand movements from local field potentials in monkey motor cortex. Nature neuroscience 6, 1253–1254, doi:10.1038/nn1158 (2003).

24 Pistohl, T., Ball, T., Schulze-Bonhage, A., Aertsen, A. & Mehring, C. Prediction of arm movement trajectories from ECoG-recordings in humans. Journal of neuroscience methods 167, 105–114, doi:10.1016/j.jneumeth.2007.10.001 (2008).

25 Rickert, J. et al. Encoding of movement direction in different frequency ranges of motor cortical local field potentials. The Journal of neuroscience : the official journal of the Society for Neuroscience 25, 8815–8824, doi:10.1523/jneurosci.0816-05.2005 (2005).

26 Schalk, G. et al. Decoding two-dimensional movement trajectories using electrocorticographic signals in humans. Journal of neural engineering 4, 264–275, doi:10.1088/1741-2560/4/3/012 (2007).

27 Thach, W. T. Correlation of neural discharge with pattern and force of muscular activity, joint position, and direction of intended next movement in motor cortex and cerebellum. Journal of neurophysiology 41, 654–676 (1978).

28 Waldert, S. et al. Hand movement direction decoded from MEG and EEG. The Journal of neuroscience : the official journal of the Society for Neuroscience 28, 1000–1008, doi:10.1523/jneurosci.5171-07.2008 (2008).

29 Witte, M., Galán, F., Waldert, S., Braun, C. & Mehring, C. Concurrent Stable and Unstable Cortical Correlates of Human Wrist Movements. Human Brain Mapping In press (2014).

30 Carmena, J. M. et al. Learning to control a brain-machine interface for reaching and grasping by primates. PLoS biology 1, E42, doi:10.1371/journal.pbio.0000042 (2003).

31 Ethier, C., Oby, E. R., Bauman, M. J. & Miller, L. E. Restoration of grasp following paralysis through brain-controlled stimulation of muscles. Nature 485, 368–371, doi:10.1038/nature10987 (2012).

32 Moritz, C. T., Perlmutter, S. I. & Fetz, E. E. Direct control of paralysed muscles by cortical neurons. Nature 456, 639–642, doi:10.1038/nature07418 (2008).

33 Serruya, M. D., Hatsopoulos, N. G., Paninski, L., Fellows, M. R. & Donoghue, J. P. Instant neural control of a movement signal. Nature 416, 141–142, doi:10.1038/416141a (2002).

34 Taylor, D. M., Tillery, S. I. & Schwartz, A. B. Direct cortical control of 3D neuroprosthetic devices. Science (New York, N.Y.) 296, 1829–1832, doi:10.1126/science.1070291 (2002).

35 Velliste, M., Perel, S., Spalding, M. C., Whitford, A. S. & Schwartz, A. B. Cortical control of a prosthetic arm for self-feeding. Nature 453, 1098–1101, doi:10.1038/nature06996 (2008).

36 Ghez, C., Gordon, J. & Ghilardi, M. F. Impairments of reaching movements in patients without proprioception. II. Effects of visual information on accuracy. Journal of neurophysiology 73, 361–372 (1995).

37 Gordon, J., Ghilardi, M. F. & Ghez, C. Impairments of reaching movements in patients without proprioception. I. Spatial errors. Journal of neurophysiology 73, 347–360 (1995).

38 Rothwell, J. C. et al. Manual motor performance in a deafferented man. Brain : a journal of neurology 105 **(Pt** **3**), 515–542 (1982).

39 Sanes, J. N., Mauritz, K. H., Evarts, E. V., Dalakas, M. C. & Chu, A. Motor deficits in patients with large-fiber sensory neuropathy. Proceedings of the National Academy of Sciences of the United States of America 81, 979–982 (1984).

40 Wolpert, D. M., Ghahramani, Z. & Jordan, M. I. An internal model for sensorimotor integration. Science (New York, N.Y.) 269, 1880–1882 (1995).

41 Wolpert, D. M. & Miall, R. C. Forward Models for Physiological Motor Control. Neural networks : the official journal of the International Neural Network Society 9, 1265–1279 (1996).

42 Guillot, A., Di Rienzo, F., MacIntyre, T., Moran, A. & Collet, C. Imagining is not doing but involves specific motor commands: a review of experimental data related to motor inhibition. Frontiers in human neuroscience 6, doi:10.3389/fnhum.2012.00247 (2012).

43 Miller, K. J. et al. Cortical activity during motor execution, motor imagery, and imagery-based online feedback. Proceedings of the National Academy of Sciences of the United States of America 107, 4430–4435, doi:10.1073/pnas.0913697107 (2010).

44 Wang, W. et al. Decoding and cortical source localization for intended movement direction with MEG. Journal of neurophysiology 104, 2451–2461, doi:10.1152/jn.00239.2010 (2010).

45 Pandarinath, C., Gilja, V., Blabe, C. H., Hochberg, L. R., Shenoy, K. V. & Henderson J. M. Overlapping neural representations of upper extremity movements in human primary motor cortex during volitional, imagined, observed, and passive movements. Program No. 80.03. 2013 Neuroscience Meeting Planner. Washington, DC: Society for Neuroscience. Online. San Diego, CA. (2013).

46 Herter, T. M., Korbel, T. & Scott, S. H. Comparison of neural responses in primary motor cortex to transient and continuous loads during posture. Journal of neurophysiology 101, 150–163, doi:10.1152/jn.90230.2008 (2009).

47 Suminski, A. J., Tkach, D. C., Fagg, A. H. & Hatsopoulos, N. G. Incorporating feedback from multiple sensory modalities enhances brain-machine interface control. The Journal of neuroscience : the official journal of the Society for Neuroscience 30, 16777–16787, doi:10.1523/jneurosci.3967-10.2010 (2010).

48 Gaunt, R. A., Collinger, J. L., Wodlinger, B., Weber, D. J. & Boninger, M. L. Propioceptive feedback enables brain computer interface (BCI) controlled prosthetic arm movement in the absence of visual input. Program No. 374.12. 2013 Neuroscience Meeting Planner. Washington, DC: Society for Neuroscience. Online. San Diego, CA. (2013).

49 Scott, S. H. The computational and neural basis of voluntary motor control and planning. Trends in cognitive sciences 16, 541–549, doi:10.1016/j.tics.2012.09.008 (2012).

50 Todorov, E. & Jordan, M. I. Optimal feedback control as a theory of motor coordination. Nature neuroscience 5, 1226–1235, doi:10.1038/nn963 (2002).

51 Wolpaw, J. R., Birbaumer, N., McFarland, D. J., Pfurtscheller, G. & Vaughan, T. M. Brain–computer interfaces for communication and control. Clinical Neurophysiology 113, 767–791, doi:http://dx.doi.org/10.1016/S1388-2457(02)00057-3 (2002).

52 Fetz, E. E. Volitional control of neural activity: implications for brain–computer interfaces. The Journal of physiology 579, 571–579, doi:10.1113/jphysiol.2006.127142 (2007).

53 Birbaumer, N., Murguialday, A. R. & Cohen, L. Brain-computer interface in paralysis. Current opinion in neurology 21, 634–638, doi:10.1097/WCO.0b013e328315ee2d (2008).

54 Cole, J. Pride and a Daily Marathon. (MIT Press, 1995).

55 Baillet, S., Mosher, J. C. & Leahy, R. M. Electromagnetic brain mapping. Signal Processing Magazine, IEEE 18, 14–30, doi:10.1109/79.962275 (2001).

56 Gramfort, A., Papadopoulo, T., Olivi, E. & Clerc, M. OpenMEEG: opensource software for quasistatic bioelectromagnetics. Biomedical engineering online 9, 45, doi:10.1186/1475-925x-9-45 (2010).

57 Holmes, C. J. et al. Enhancement of MR images using registration for signal averaging. Journal of computer assisted tomography 22, 324–333 (1998).

58 Alexandre, L. A., Campilho, A. C. & Kamel, M. On combining classifiers using sum and product rules. Pattern Recognition Letters 22, 1283–1289, doi:http://dx.doi.org/10.1016/S0167-8655(01)00073-3 (2001).

59 Fukunaga, K. Introduction to statistical pattern recognition. (Academic Press, 1972).

60 Tadel, F., Baillet, S., Mosher, J. C., Pantazis, D. & Leahy, R. M. Brainstorm: a user-friendly application for MEG/EEG analysis. Computational intelligence and neuroscience 2011, 879716, doi:10.1155/2011/879716 (2011).

